# Prevalence and patterns of higher-order interactions

**DOI:** 10.1101/233312

**Authors:** Elif Tekin, Cynthia White, Tina Manzhu Kang, Nina Singh, Mauricio Cruz-Loya, Robert Damoiseaux, Van M. Savage, Pamela J. Yeh

**Affiliations:** Department of Ecology and Evolutionary Biology, University of California, Los Angeles, CA 90095; Department of Biomathematics, University of California, David Geffen School of Medicine, Los Angeles, CA 90095; Department of Medical and Molecular Pharmacology, University of California, David Geffen School of Medicine, Los Angeles, CA 90095; Santa Fe Institute, Santa Fe, NM 87501

**Keywords:** higher-order interactions, emergent interactions, antibiotic

## Abstract

Interactions and emergent processes are essential for research on complex systems involving many components. Most studies focus solely on pairwise interactions and ignore higher-order interactions among three or more components. To gain deeper insights into higher-order interactions and complex environments, we study antibiotic combinations applied to pathogenic *Escherichia coli* and obtain unprecedented amounts of detailed data (251 two-drug combinations, 1512 three-drug combinations, 5670 four-drug combinations, and 13608 five-drug combinations). Directly opposite to previous assumptions and reports, we find higher-order interactions increase in frequency with the number of drugs in the bacteria’s environment. Furthermore, we observe a shift towards net synergy (effect greater than expected based on independent individual effects) and towards emergent antagonism (effect less than expected based on lower-order interaction effects). These findings have implications for the potential efficacy of drug combinations and are crucial for better navigating problems associated with the combinatorial complexity of multi-component systems.

## Introduction

Interactions are the key to unlocking emergent and unintuitive properties across many fields: reactions in biochemistry, food webs and flocking among birds in ecology, environmental stressors and effects on species diversity in conservation biology, genetic interactions in evolution and bioinformatics, bound states and many-body interactions in physics, social interactions in economics and political science, and drug interactions in pharmacology ^1, 2, 3, 4, 5, 6^. Understanding whether components interact in a manner that enhances (synergy) or weakens (antagonism) the individual effects of the mixed components is important because the type of interaction governs the dynamics of complex systems. For example, in conservation biology, understanding multiple-stressor effects informs the development of strategies to prevent loss of biodiversity ^6^. In pharmacology, understanding drug interactions enables the effective design of treatment strategies to combat complex diseases such as cancer ^7^ and HIV ^8^, which increasingly rely on multi-drug treatments. Despite the structural and terminological differences between natural and social systems, multi-component interactions constitute a unifying theme in studying and predicting patterns in large complex systems.

The formation of complex structures and dynamics often results from emergent properties that cannot be explained based on the effects of individual components or even interactions between pairwise parts or other lower-order interactions (fewer numbers of components than the whole combination). For instance, addition of a small amount of a third drug may alter the interaction between two drugs already being used. Moreover, some interactions only exist when many components are present, even though there is no interaction between any of the isolated pairs or triples ^9^, such as a protein or molecule that requires all parts to be joined before it can properly function and any activity or response can be measured.

Therefore, an approach that explicitly distinguishes emergent interactions—interactions that are not due to the presence of lower-order interactions—from the existence of any (net) interaction is indispensable to fully represent complex system dynamics ^10, 11^.

Despite the possibility of higher-order interactions, studies have often focused on pairwise interactions ^12, 13, 14, 15^ while presuming or concluding that higher-order effects—due to three or more components—are either extremely rare or negligible ^16, 17, 18^. There are both conceptual and practical reasons that have led to this view. On the conceptual side, it has been commonly argued that lower-order effects are likely to counteract each other in higher-order combinations such that they essentially cancel out and result in zero or negligible net effect ^19, 20^. Alternatively, it has been argued that higher-order interactions prevent large systems from exhibiting stability, and thus concluded that these effects either do not exist or are insignificant ^21^. For these reasons, many studies either implicitly assume higher-order interactions do not exist or argue against the importance of measuring and considering higher-order interactions ^13, 16, 22, 23^.

A practical reason why higher-order interactions have received much less attention relative to two-way (pairwise) interactions is due to combinatorial complexity, i.e., difficulty in collecting data for all subsets of combinations of components ^15,16^. Another practical challenge to examining and classifying higher-order interactions is that it entails calculating the contribution from all lower levels—subsets of fewer components—to the emergent behavior of the whole combination. These calculations are surprisingly subtle and correspond to having well-defined and understood emergent interaction metrics with straightforward generalization to higher orders and quantification of uncertainties. The lack of such methods in many fields has stood as a theoretical limitation for studying interacting systems ^10, 14^.

Recent progress has made it possible to overcome these latter two practical limitations and thus enables us to directly test the above conceptual presumption by measuring how frequent higher-order interactions are and what types of interactions are present. In this regard, our recent studies of combinations of three stressors ^9, 24^ showed that many more interactions arise with three-way interactions compared with two-way interactions. Moreover, these three-way interactions exhibit an important feature: higher degrees of emergent antagonism, compared with pairwise interactions. This greater amount of emergent antagonism—reduced effect relative to the expected effect based on no-interaction—suggests a need to further explore interactions among three, four, and five drugs. Correspondingly, an insightful study by Mayfield et al. ^13^ found that incorporating higher-order interactions as opposed to a constrained approach of two-way interactions leads to better expectations of variations in natural plant communities, and a very recent and intriguing review by Levine et al. ^14^ has provided a theoretical approach that shows how higher-order interactions can promote species richness. In addition, another compelling study demonstrated the existence of higher-order gene interactions and revealed how they can explain complex alterations in gene expression ^25^.

These studies have shown the presence of higher-order effects and the importance of exploring them. However, what is lacking is a comprehensive framework and dataset that can be used to reveal the patterns of higher-order interactions. In particular, a feasible empirical study design (which provides easy manipulation of the environmental disturbances and the number of components in the system) and a comprehensive theoretical approach are necessary for gaining crucial insights into biological systems (or more generally, complex systems) comprised of many components. Indeed, the theoretical framework in this setting should be conveniently generalizable to distinguish different types of higher-order effects (i.e., net versus emergent) in the presence of any number of components.

Using large numbers of drug combinations, we ask: 1) Are there more (or fewer) interactions when more stressors are added? 2) Are there any trends in the frequency of interaction types that amplify or attenuate the complexity of a system when the number of components is increased? 3) Do these trends in interaction type differ between net versus emergent interactions?

In contrast to previous work, we show that emergent higher-order interactions are much more pervasive than commonly assumed and that there are characteristic patterns of net versus emergent higher-order interactions as the number of drugs increases. Addressing these issues will be extremely useful if we are to find a perspicuous path forward in searching for higherorder interactions and dealing with combinatorial complexities.

In this paper, we ground these questions in a highly-controlled antibiotic study system that examines bacterial growth responses to an environment that consists of different drug combinations. We obtain data for how 8 single drugs with 3 concentrations each, 251 two-drug combinations, 1512 three-drug combinations, 5670 four-drug combinations, and 13608 five-drug combinations affect pathogenic *E. coli* growth rates (Fig 1). This full factorial design of drug combinations allows characterization of net and emergent interactions for all five-way and lower-order interactions (two-, three-, four-way), and thus represents a staggering amount of data compared with previous studies, allowing us to shed new light on how interactions change as more and more drugs (components) are added.

**Fig 1.**
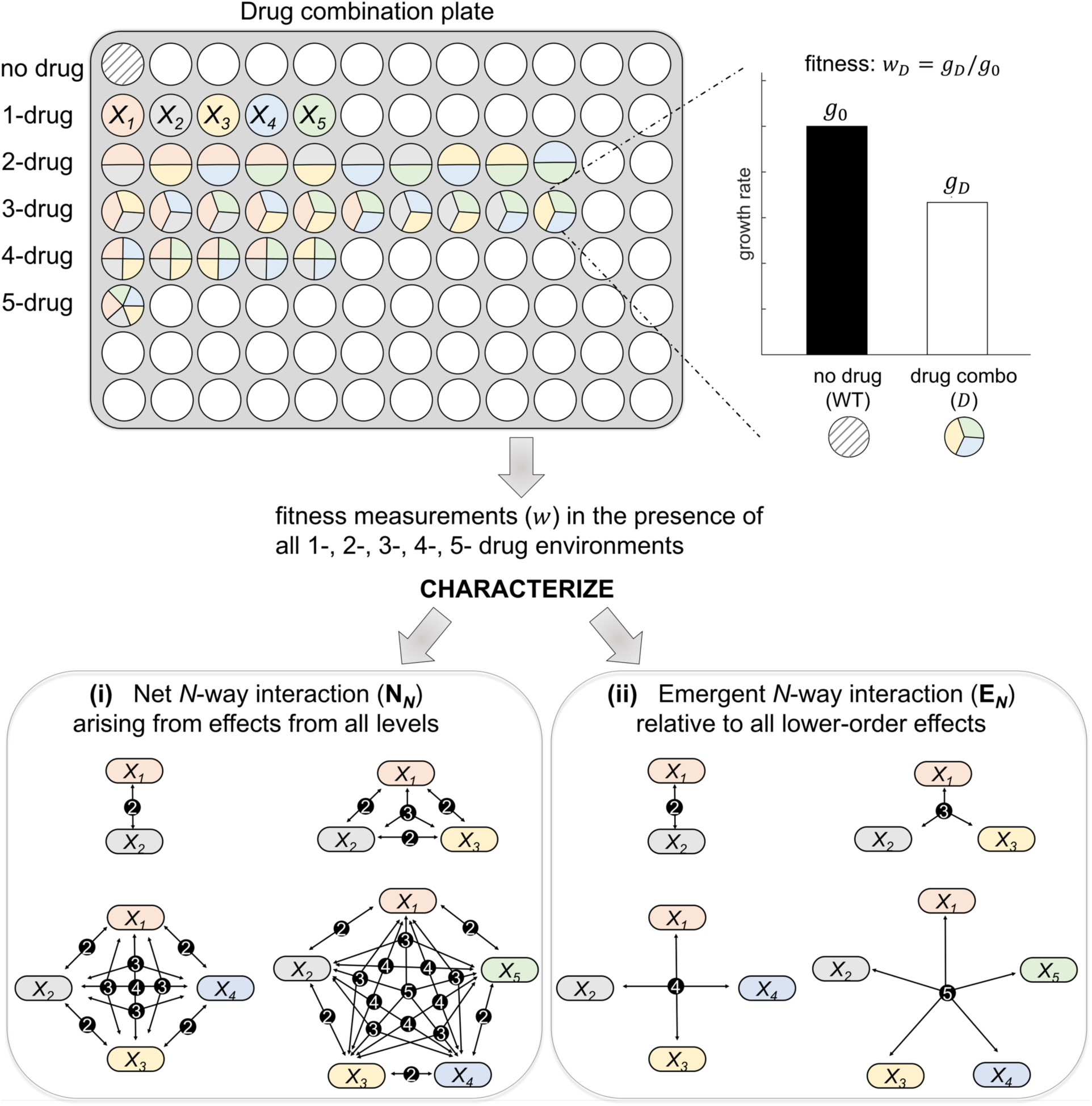
Experimental and theoretical setup for the characterization of higher-order interactions. Schematic representation of a drug-combination plate, where the shaded well in the first row represents the control strain with no drug added and colored wells correspond to single or *N*-way (up to 5-way) combinations from a set of drugs denoted by *X_i_*. The N-way combinations of drugs are represented by wells divided into *N* identical slices with the colors signifying the drugs in the combination (see 1-drug row for each color). The concentration of each drug is kept the same across single, 2-, 3-, 4-, and 5-drug combination experiments. Here, the experimental setup is simplified for illustration, but in actuality, 1) we filled all wells with bacteria and drug combinations to obtain replicate measurements (see “Experimental details”), and 2) we used multiple 96-well plates for each 5-drug combination. From these experiments, fitness of bacteria (*w*) in the presence of drug combination (*D*) is assessed by the relative growth rate with respect to the no-drug control (WT). In the figure, schematics for the net N-way interaction include all possible lower-order connections, whereas an emergent interaction schematic connects all *N* drugs (such as dyad and triad for 2- and 3-drug combinations, respectively).

### Theoretical Framework for the Characterization of Higher-order Interactions

To measure interactions, we must first carefully define what an interaction is and what response measurements we use to assess the interactions. Based on standard definitions within the field, we can then extend and generalize this interactions framework to higher orders for both net and emergent interactions.

For drug studies the key response measurement is growth rate of a bacteria population in the presence of a drug relative to growth of bacteria in a no-drug environment. This growth rate is interpreted as the fitness in the presence of a drug treatment *D* and is typically denoted by *w_D_* (Fig 1), where no-growth (*w_D_* = 0) represents complete lethality and maximum-growth (*w_D_* = 1) represents the case that the drug treatment is not effective at all. Interaction classifications are then defined relative to this baseline of no-interaction (additivity) based on whether a drug combination yields more than (synergistic) or less than (antagonistic) this baseline, no-interaction (additive) effect ^10, 26^.

Here, we review net and emergent measures for the quantification of *N*-way interaction effects (see Fig 1 and S1 Fig). Throughout the paper, we denote single drugs by *X_i_*, where *i* >0 and use a list of *X_i_* to represent drug combinations (such as *X*_1_*X*_2_ for combining two drugs *X*_1_ and *X*_2_). Moreover, for notational tractability, N-way interaction measures are defined for *X*_1_*X*_2_…*X_N_*, which stands for any *N*-drug combination.

#### Net *N*-way (N_*N*_) interactions

We follow the Bliss Independence model ^27^ to characterize a net *N*-way interaction (N_*N*_) for the total interaction in comparison to that expected from all the independent and individual effects of each drug. Based on Bliss Independence, drugs *X*_1_ and *X*_2_ are not interacting when the addition of a second drug (*X*_2_) does not influence the percent decrease in the pathogen fitness due to another single drug (*X*_1_), i.e. *w*_*X*_1_*X*_2__ = *w*_*X*_1_*w X*_2__. Accordingly, the interaction between *X*_1_ and *X*_2_ is measured as

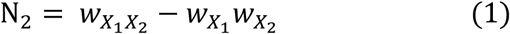

A sufficiently large negative value of N_2_ suggests a synergistic interaction, as drugs together produce a superior inhibition effect relative to the case that drugs are not interacting, whereas a large positive value of N_2_ indicates an antagonistic interaction. However, a correct rescaling method ^28^ is needed to properly interpret the magnitude of these measures.

Extending Eq. (1) to systems with more than two drugs enables measurements and calculations to determine the presence of any kind of interaction relative to the single-drug effects. Thus, the generalized form of the net interaction measure for an *N*-drug combination is ^10, 29, 30^.

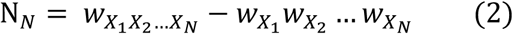

Observe that the subscript *X*_1_*X*_2_…*X_N_*means the bacteria’s environment contains drug *X*_1_ plus drug *X*_2_ plus all drugs up to *X_N_*. Therefore, if only *k* of the *N* drugs remain in the environment, the subscript becomes *X*_1_*X*_2_…*X_k_*0…0, which is equivalent to *X*_1_*X*_2_…*X_k_*because adding 0 or no drug is equivalent to just having the *k*-drug subset. Moreover, the relative fitness of the drugs at 0 concentration will be 1, so the net *N*-way interaction will reduce to just a *k*-way interaction, as expected.

#### Emergent *N*-way (E_*N*_) interactions

To assess interactions that require all *N* drugs to be present, or equivalently, interactions beyond what is expected from the effects of all lower-order combinations, we use and extend our emergent interaction framework presented in Beppler et al. ^9^. When two drugs are combined, the definitions of net and emergent interactions converge to become identical because single drugs constitute the one and only lower-order component of a two-drug environment, so there is nothing from which to emerge except the single-order effects. Thus, the emergent 2-way (E_2_) interaction is identical to the net 2-way interaction, N_2_ (Eq. (1)). However, when there are more than two drugs in the environment, interactions among different lower-order subsets (2-way versus 3-way versus 4-way or the combination of 2-ways, etc.) of drugs can change the dynamics of N-way interactions, and those effects need to be accounted for by characterizing the emergent interactions ^11^. Here, the contribution to the effect that comes solely from a lower-order combination corresponds to the total interaction when only that specific lower-order interaction is present. Based on this, we systematically determine all the lower-order parts in an *N*-way combination and subtract off those effects from the net interaction to capture any emergent interaction (see details in the Materials and Methods). When *N*=3, this corresponds to the total (net) three-drug interaction effect that is not due to the contributions from all the two-drug combinations. As described in Materials and Methods, E_3_ yields an expression that includes fitness measurements in the presence of every possible drug combination in the three-drug environment.

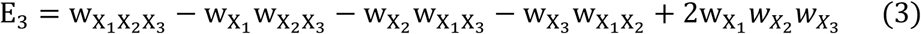

The emergent 4-way and 5-way interaction formulas are derived in the Materials and Methods with explanation given for how to correctly count the combinatorics of all drug subsets while avoiding any issues of double- or over-counting of contributions.

## Results

In this study, we explored the consequences of increased complexity in multi-component systems by employing an experimental design of higher-order drug combinations and by using and extending our recently developed mathematical framework ^9, 24^ to characterize higher-order interactions. We categorized net interactions based on the deviation of the whole combination effect from the expected effect of no-interaction among individual drugs. To measure emergent interactions, we calculated the deviation of net interaction from the expected effect from all interactions that result from lower-order subsets/combinations of drugs (Fig 1, S1 Fig, “Theoretical Framework for the Characterization of Higher-order Interactions” and “Materials and Methods”). As with pairwise studies for drugs, epistasis, and other biological systems, interactions are defined as synergistic or antagonistic when the effect of two components is sufficiently (see Materials and Methods for precise values) stronger or weaker than expected effects of no-interaction (net) or all lower-order interactions (emergent), respectively.

As shown in Fig 2A, (No-interaction bar) and S2 Fig, interactions become significantly more frequent as the number of drugs in *E. colf*’s environment increases (sum of Synergy and Antagonism bars in Fig 2A). This startling finding suggests not only that higher-order interactions are not negligible and should not be ignored, but that they may be even more important than pairwise interactions in determining the structure and dynamics of systems because they are substantially more prevalent than two-component interactions. Understanding this phenomenon is thus fundamental to understanding interactions in biological and complex systems in general. Evaluating whether any patterns exist in the types of interactions as the number of drugs increases, we found that the net interactions among drugs tend towards more synergy, whereas emergent interactions exhibit a shift towards more antagonism (Fig 2).

**Fig 2.**
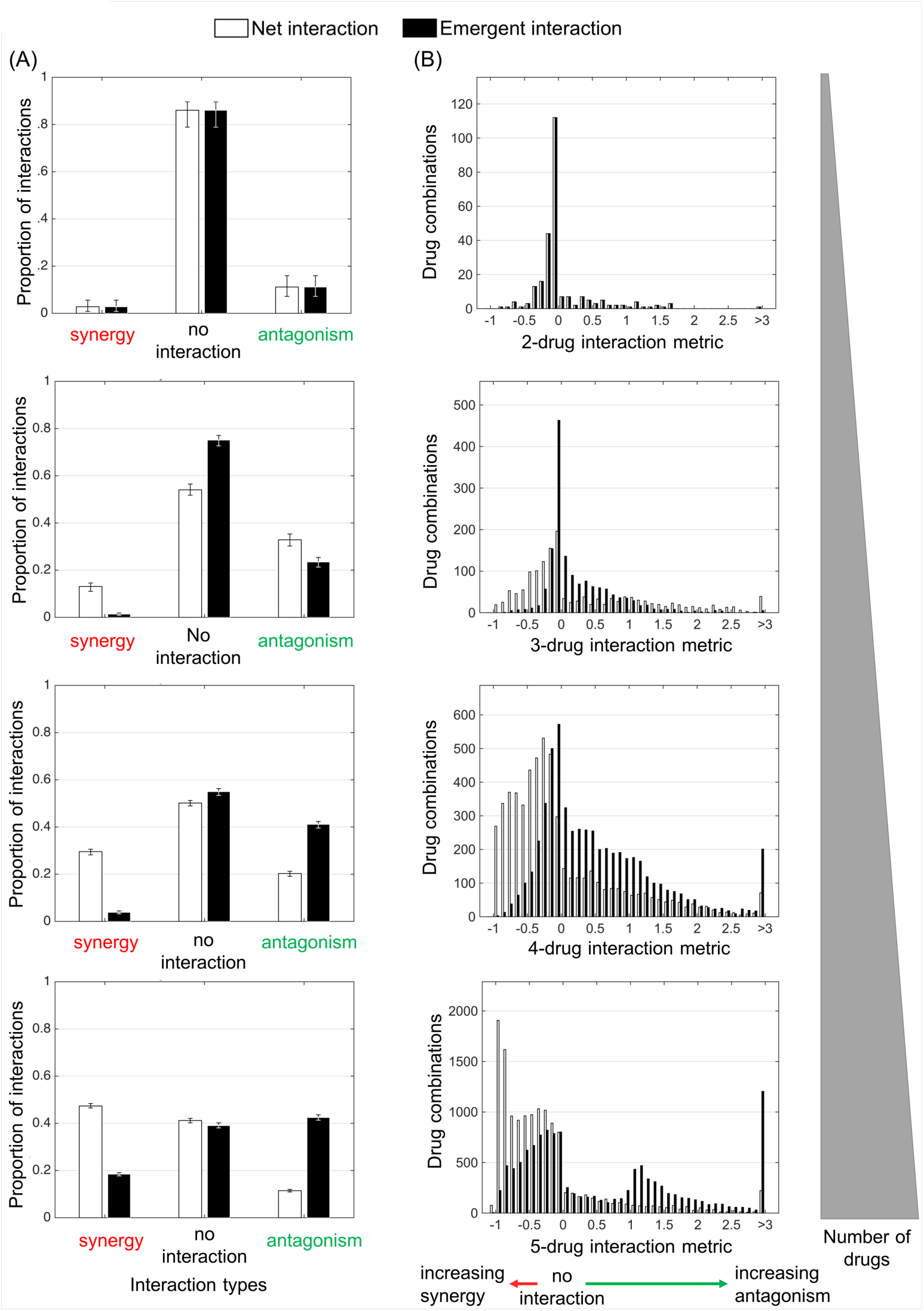
Overall behavior of interaction metric results and categorization of interactions. For each *N*-way interaction, (A) bar plots for interaction classifications (synergy, no-interaction, antagonism) of net (white) and emergent (black) interactions with 95% confidence intervals resulting from bootstrapping experiments via sampling with replacement over all measured drug combinations and (B) histograms of net (white) and emergent (black) interaction metric results with a bin size of 0.1 are plotted. A diagram is shown that displays the direction in which the strength of specific interaction class enlarges. Note here that the definitions of net 2-way and emergent 2-way interactions are identical. Hence, plots corresponding to the distribution of interaction metrics and the proportion of interactions are indistinguishable for 2-drug combinations.

Further dissecting the nature of these interactions and comparing net with emergent interactions, we found that net synergy seldom implies emergent synergy (Fig 3A, Synergy column). This makes sense because any interactions will create a net effect while emergence is only an interaction among all drugs in the combination, so will almost certainly be a subset of net interactions. Consequently, a net three-way synergy is usually due to pairwise synergies between two drugs in which a third drug may not be increasing efficacy but still increases toxicity to patients.

**Fig 3.**
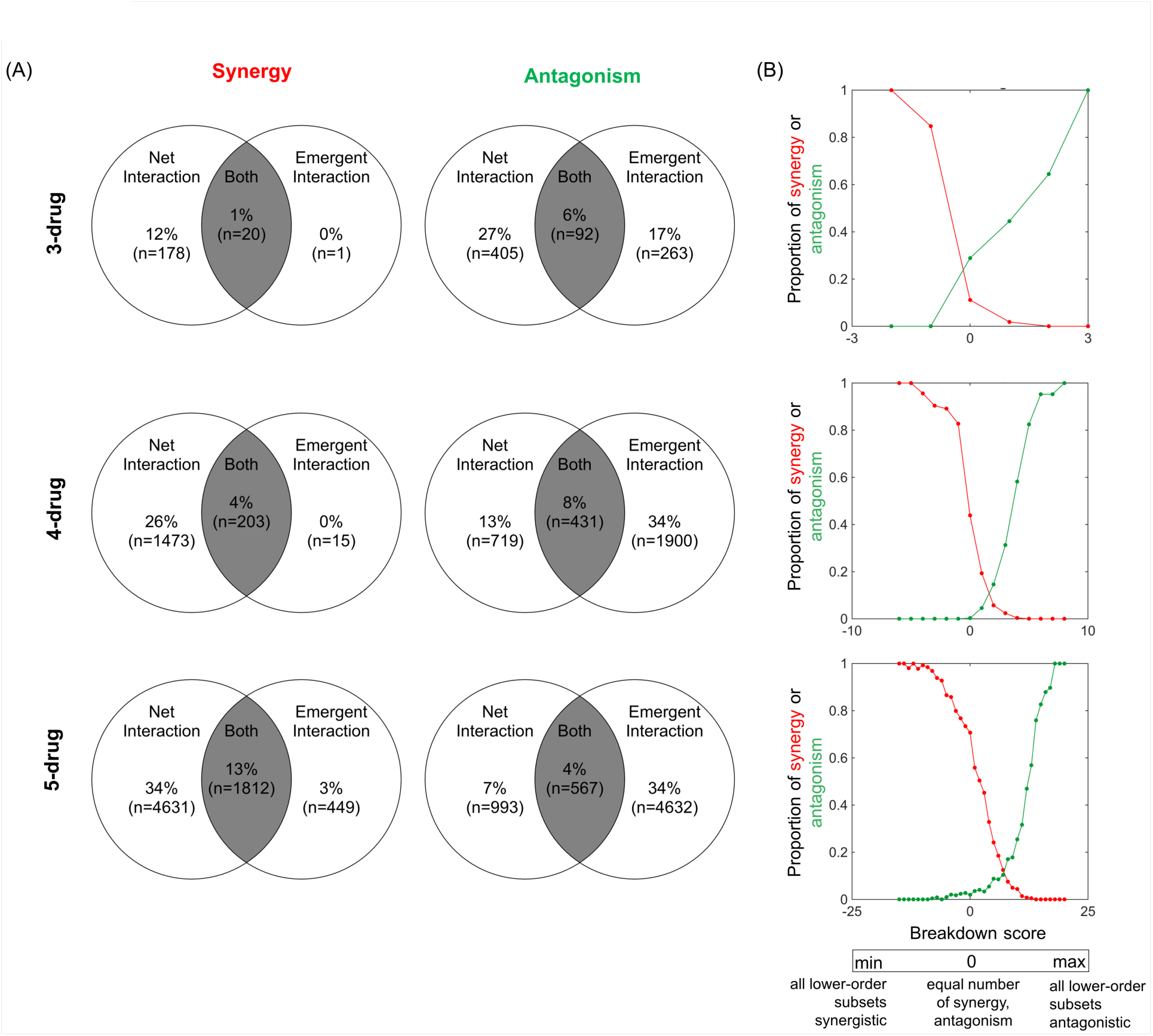
Comparison of net and emergent interactions by synergy and antagonism. (A) Venn diagrams comparing an overlap for different interaction categorizations (synergy: left column, antagonism: right column) according to net and emergent interaction measures of 3-, 4-, and 5-drug combinations are given. For each N-drug Venn diagram, the percentages are calculated relative to the number of *N*-drug combinations. (B) The proportion of net synergies and net antagonisms are plotted versus the total breakdown score. Breakdown scores represent the dominant form of interaction types at the lower-order combinations calculated by the summation of −1, and 1 over each lower-order synergy and lower-order antagonism, respectively (see S1 Table). The minimum and maximum values of breakdown score differ across each plot as the total number of lower-order combinations of an N-drug combination depends on the value of *N*.

In addition, we found that emergent antagonism is less likely to imply net antagonism as the number of drugs increases (Fig 3A, Antagonism column). Explicitly, this fraction is given by 26%, 18%, and 11% at the 3-, 4-, and 5-drug level, respectively. Intriguingly, we showed that lower-order net synergistic effects overcome lower-order net antagonistic effects (Fig 3B). This demonstrates the fallacy of the prominent presumption that higher-order interactions cancel out and are negligible. Establishing this result was only possible due to our full-factorial experiments and large-scale dataset as well as our development and comparison of emergent versus net interaction measures. Overall, our comparison analysis of net and emergent interactions indicates that emergent synergy mostly suggests net synergy, whereas emergent antagonism does not imply net antagonism.

## Discussion

We have explored the consequences of increasing the number of components in the context of net and emergent higher-order interactions through a systematic analysis of bacterial responses in the presence of drug combinations. For both net and emergent higher-order interactions we found the number of interactions substantially increased as the number of drugs increased. Although this may not be surprising for net interactions, *it is extremely surprising for emergent interactions because these have been largely ignored in the literature, yet our new analysis reveals emergent interactions are highly prevalent in our drug systems*. Although we have only shown this result for drug combinations, it contradicts the prevailing views and previous limited results in the field.

We also observed an increasing trend towards synergy in net interaction effects and towards antagonism in emergent interaction effects. These trends use tremendous amounts of data to extend and elaborate on recent patterns found in the comparison of two- with three-drug combinations ^24^ These general trends suggest that increasing the number of drugs continually adds new layers of complexity and leads to the natural question of whether this layering of complexity also applies to interactions among multiple components across a myriad of other systems. Indeed, if this finding continues to hold as the number of drugs increases and also applies to other systems and fields, this bypass approach could revolutionize studies of both net and emergent higher-order interactions by making the intractability of the combinatorics suddenly become tractable via systematic patterns that enable predictability.

From a clinical standpoint, synergies offer higher treatment efficacies with low toxicity; hence, they are valued and used clinically while antagonistic combinations have been traditionally avoided ^11^. Our study reveals for the first time, to our knowledge, that for most higher-order drug combinations, net synergy does not imply emergent synergy. The abundance of such cases suggests the criteria for determining clinically advantageous (or disadvantageous) drug combinations should now consider both net and emergent effects as it is also critical to identify whether addition of drugs yields a real (emergent) benefit that justifies the inclusion of each additional drug in the combination.

Studies on pairwise-drug combinations have shown that antagonistic interactions can lead to a selective advantage of the wild-type pathogen population and reduce the rate of adaptation to drugs ^31, 32^. Extending these ideas to more rugged fitness landscapes that correspond to higherorder interactions among drugs and determining the consequences for drug-resistance has not yet been pursued. To our knowledge, this is one of the only and by far the largest set of empirical data obtained to examine the role of net and emergent higher-order interactions. Our observations suggest that the fitness landscapes for multi-drug combinations should be extremely rugged due to the pervasiveness of higher-order interactions.

Although our study focuses on a drug-bacteria system, the conceptualization of interaction types (net versus emergent, and synergy versus antagonism) and the systematic analysis of higher-order interactions can be extrapolated into other fields. A relatively straightforward application of our framework includes multiple stressor effects since drugs in our study are essentially stressors to the bacterial population. Moreover, the interaction model used in multiple predator effect studies is equivalent to the net interaction measure ^33^, suggesting a strong correspondence of our mathematical framework for characterizing emergent interactions ^9^. In microeconomics, relationships between individuals (creating demand) and market firms (supplying demand) affect the allocation of resources ^2^. In the context of social groups, the idea of emergence can be well represented by one individual’s role in controlling a conflict between others in the group. Indeed, studies in primates ^34^ and in people ^35^ have predicted that cohesion dynamics of groups differ significantly between dyads and triads—groups of two and three people, respectively—with triads being more stable in conserving the association of a group.

Here, we also note several caveats in our framework in its application to clinical practice as well as to other interaction-network settings. From a clinical standpoint, drug combination experiments in this study are carried out *in vitro*, in highly controllable systems, hence *in vivo* studies of higher-order drug combinations are needed for guiding studies with clinical applications. In addition, as opposed to the drug-bacteria system, natural systems are comprised of many interacting species and functional groups that span trophic levels and that are impacted by diverse but sometimes correlated environmental drivers such as temperature, precipitation, stoichiometry, fires, etc. Consequently, new theory needs to be developed that incorporates additional information to study higher-order interactions and their outcomes in natural systems that involve many more complexities.

In conclusion, we introduce an approach to studying higher-order interactions in biological systems. We provide an enormous amount of data that we analyze with a recently developed theoretical framework to characterize net and emergent interactions among all possible combinations of 2, 3, 4, and 5 drugs out of a set of 8 antibiotics. Notably, we find many interactions that only emerge when multiple drugs are present, and even more surprisingly, we find that the frequency of interactions increases as the number of components in the system increases, contradicting the assumptions and limited findings of many previous studies. Beyond this, we find that emergent interactions tend towards antagonism while net interactions tend towards synergy. These findings suggest that higher-order interactions may be of fundamental importance in understanding and predicting the structure and dynamics of complex biological systems with many interacting parts, such as drugs, genes, food webs, environmental stressors, and more.

Intriguingly, the general questions explored here for drug interactions are highly relevant for other fields, and we anticipate that the research program presented here will be useful for revealing higher-order emergent properties and patterns in ecological, medical, evolutionary, and social systems. It is possible that the prevalence and patterns of higher-order interactions described here will be generic to many other biological and complex systems. Alternatively, these findings may be specific to the type of system being studied. The answer can only be elucidated through future work in other systems. For our drug-bacteria system, we have combined a large-scale experimental system with a new conceptual framework to establish the strong prevalence and importance of higher-order interactions and to identify patterns of these interactions that should help to circumvent combinatorial complexity as well as to inform effective design of multi-drug treatments.

## Materials and Methods

### Experimental details

#### Bacterial storage and preparation

Pathogenic *E. Coli* strain CFT073 (ATCC^®^ 700928™) isolated from human clinical specimens was used for all experiments. Bacterial aliquots were made using a single colony collected using streak purification. The aliquots were stored in 25% glycerol at −80°C. A culture was prepared each day using a thawed aliquot diluted 10^−2^ in Luria Broth (10 g/l tryptone, 5 g/l yeast extract, and 10 g/l NaCl). The culture was grown at 37°C for about 4 hours.

#### Antibiotics

We used eight antibiotics in this study that were chosen specifically to cover a broad range of antibiotic mechanisms of action ^36^. Additionally, drugs needed to be soluble in DMSO and therefore were chosen based on solubility properties. These antibiotics were Ampicillin (Sigma A9518), Cefoxitin Sodium Salt (Sigma C4786), Ciprofloxacin Hydrochloride (MP Biomedicals 199020), Doxycycline hyclate (Sigma D9891), Erythromycin (Sigma Aldrich E6376), Fusidic Acid Sodium Salt (Sigma F0881), Streptomycin (Sigma Aldrich S6501), and Trimethoprim (Sigma T7883). A list of drugs, abbreviations, mechanisms of action, and concentrations used in this assay is in Table 1.

**Table 1.**
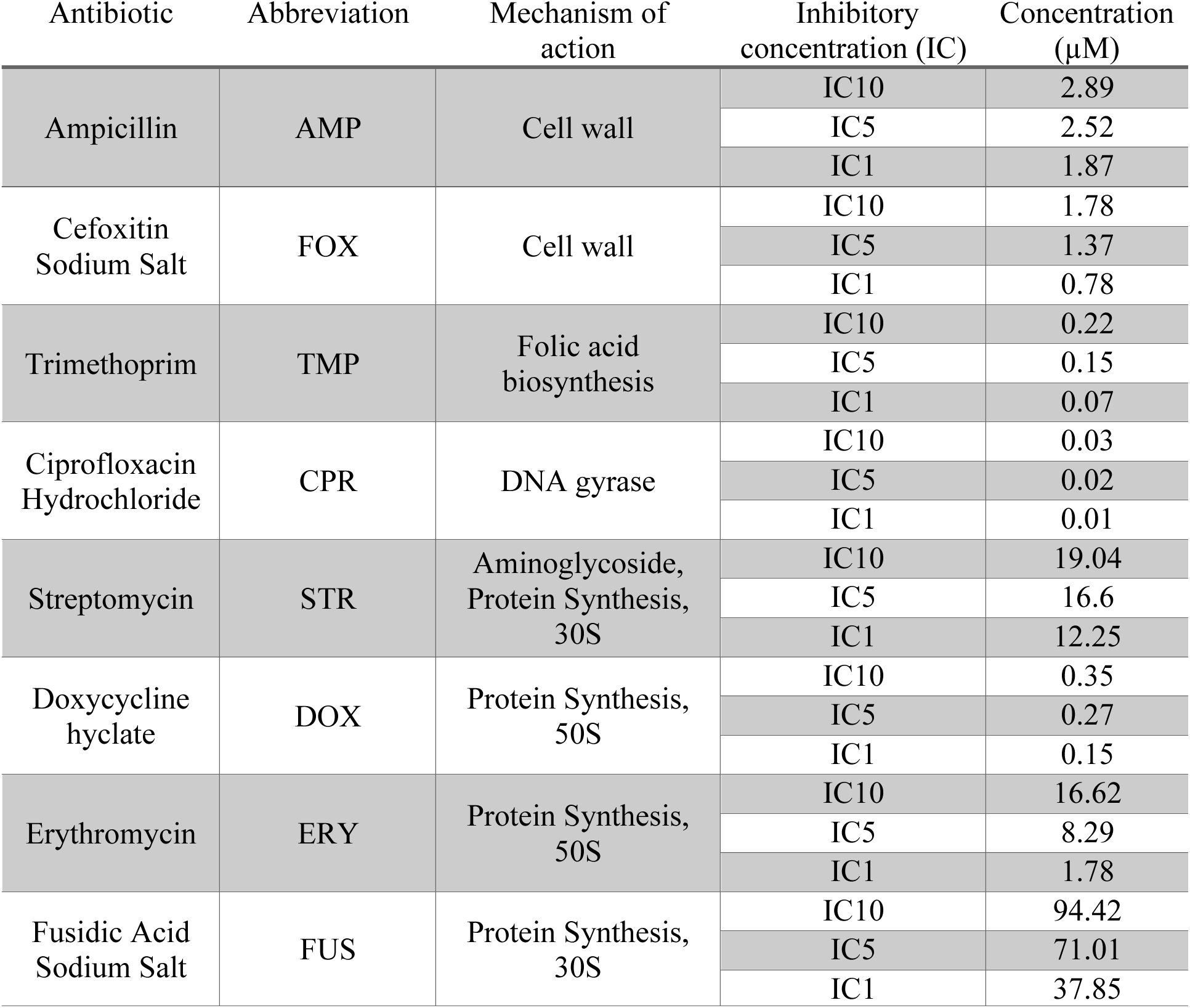
Summary of antibiotics.

| Antibiotic | Abbreviation | Mechanism of action | Inhibitory concentration (IC) | Concentration (μM) |
| --- | --- | --- | --- | --- |
| Ampicillin | AMP | Cell wall | IC10 | 2.89 |
|  |  |  | IC5 | 2.52 |
|  |  |  | IC1 | 1.87 |
| Cefoxitin Sodium Salt | FOX | Cell wall | IC10 | 1.78 |
|  |  |  | IC5 | 1.37 |
|  |  |  | IC1 | 0.78 |
| Trimethoprim | TMP | Folic acid biosynthesis | IC10 | 0.22 |
|  |  |  | IC5 | 0.15 |
|  |  |  | IC1 | 0.07 |
| Ciprofloxacin Hydrochloride | CPR | DNA gyrase | IC10 | 0.03 |
|  |  |  | IC5 | 0.02 |
|  |  |  | IC1 | 0.01 |
| Streptomycin | STR | Aminoglycoside, Protein Synthesis, 30S | IC10 | 19.04 |
|  |  |  | IC5 | 16.6 |
|  |  |  | IC1 | 12.25 |
| Doxycycline hyclate | DOX | Protein Synthesis, 50S | IC10 | 0.35 |
|  |  |  | IC5 | 0.27 |
|  |  |  | IC1 | 0.15 |
| Erythromycin | ERY | Protein Synthesis, 50S | IC10 | 16.62 |
|  |  |  | IC5 | 8.29 |
|  |  |  | IC1 | 1.78 |
| Fusidic Acid Sodium Salt | FUS | Protein Synthesis, 30S | IC10 | 94.42 |
|  |  |  | IC5 | 71.01 |
|  |  |  | IC1 | 37.85 |

The antibiotics used are listed with their mechanism of action and concentrations corresponding to 10%, 5%, and 1% inhibitory concentration levels (IC10, IC5, and IC1, respectively).

#### Antibiotic concentration determination and preparation

A dose curve was generated to determine antibiotic drug concentrations for this assay. Dose curves were generated using 20 drug concentrations with a dilution factor of two and a starting concentration of 0.1 mM. For Fusidic Acid, the highest concentration was 1 mM, as the lower concentrations were found to be ineffective in generating lethality needed to determine the IC50 (50% Inhibition Concentration). Graphpad Prism 7 was used to graph the dose curve and determine the IC50. Additionally, Graphpad (http://www.graphpad.com/quickcalcs/Ecanything1/) was used to estimate the IC10, IC5, and IC1. Minimally effective concentrations were chosen in order to maintain bacterial growth even in multidrug combinations. Each antibiotic was weighed and solubilized in 100% DMSO (Sigma), except for Streptomycin which was solubilized in 50% DMSO, to a final concentration of 400x the determined IC10, IC5, and IC1 concentrations. Antibiotics were then pipetted into a source plate (Thermo Scientific) as either a single antibiotic with additional DMSO or in combination with another antibiotic. The final concentration in the source plate was 200x the target concentration.

#### Experimental setup

In total, we tested all 2-, 3-, 4-, and 5-drug combinations from a set of 8 antibiotics (Table 1), meaning that 28, 56, 70, and 56 distinct drug combinations were tested at given concentrations, respectively. For each experiment, 25 μL of Luria Broth was added to each well of a 384 well plate (Greiner BioOne) using a Multidrop 384 (Thermo Scientific). An additional 25 μL of media was added to the media-only control. Using a Biomek FX (Beckman Coulter) with a 250 nL pin tool (V&P Scientific), we pinned 250 nL from every well of each of the three premade source plates (1 plate with 1 antibiotic and DMSO and 2 plates with 2 antibiotics in combination) into the experimental plate. A 25 μL of a 10^−4^ dilution of the overday culture was then added to each well (except for the negative control). Plates were incubated at 37°C and read using an OD_590_ measurement every 4 hours for 16 hours. Each 2-, 3-, 4- and 5-drug experiment was tested at least three times (S1 Data).

#### Growth measurements

For each plate, the Z’-factor ^37^ was calculated to determine the quality of the assay. If a Z’ value was below 0.5, the plate was not used for final analysis. For a few plates, there was one well that was more than 2 standard deviations from the average of the negative control. For these plates, that well was removed from the final Z’-factor calculation. The exponential rate of growth was determined for each experimental well and compared to the average exponential rate of growth for the no drug control to give a growth percentage. Growth percentages were used to determine interaction types based on a framework developed in ^9, 24^, as summarized below.

### Mathematical framework

Here, we derive the formula for emergent 3-way interactions and also generalize the emergent interaction measure to higher orders and any *N*-way combination. As described briefly in “Theoretical Framework for the Characterization of Higher-order Interactions”, we accomplish this by starting from the definition of Bliss Independence (i.e., N_*N*_) and by subtracting all lower-order contribution effects from N_*N*_. For any *N*, we can denote these lower-order contributions by [N_*N*_]_*D*_1_⊥*D*_2_…⊥*D_n_*_, where ⊥ represents no-interaction, and *D_i_* represents one or more drugs that is a subset of interacting components (i.e., combinations of drugs) that are composed of *n* disjoint sets of drugs with the union of all such sets equaling the total set of all drugs (i.e., *D*_1_U*D*_2_U…U *D_n_* = *X*_1_*X*_2_…*X_N_*) because all subsets must combine to produce the original *N*-drug combination such that no drug is allowed to occur in more than one subset of drugs. In other words, for any lower-order contribution term, each drug can be part of only one subset. For example, [N_3_]_*X*_1_*X*_2_⊥*X*_3__, where *D*_1_= *X*_1_*X*_2_ and *D*_2_= *X*_3_, represents the pairwise combination effect of *X*_1_*X*_2_ on the 3-way interaction, and it measures the expected three-way effect when the remaining drug *X*_3_ does not interact with the pair. This effect is calculated as

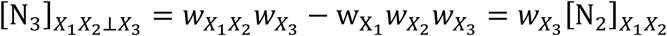

For measuring emergent interactions, all different factorizations of drugs are subtracted in a combinatorial fashion. When *N*=3, all the pairwise combination effects are subtracted from the net 3-way interaction

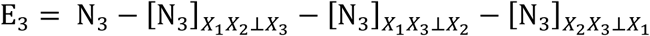

Analogously calculating the effects due to *X*_1_*X*_3_ and *X*_2_*X*_3_ (i.e., **[N_3_]_*X*_1_*X*_3_⊥*X*_2__** and **[N_3_]_*X*_2_*X*_3_⊥*X*_1__**), the definition of **E_3_** yields Eq. (3) in the “Theoretical Framework for the Characterization of Higher-order Interactions”.

The most challenging issue in deriving higher-order equations for more than three drugs is to carefully correct for the possible double counting of interactions when subtracting lower order effects that may be subsumed in multiple higher-order terms. For instance, the lower-order interaction terms of [N_4_]_*X*_1_⊥*X*_1_*X*_3_*X*_4__, and[N_4_]_*X*_2_⊥*X*_1_*X*_3_*X*_4__both subsume the effect of [N_4_]_*X*_3_*X*_4_⊥*X*_1_⊥*X*_2__ as all three are identical when *X*_1_, *X*_2_, and the combined drug pair *X*_3_*X*_4_ are all non-interacting with respect to each other. Such cases should be dealt with systematically to make sure that each lower-order effect is subtracted off exactly once (see S1 Text). Moreover, when *N* >3, it is also necessary to exclude the effect that results from interactions of different subsets of drug combinations. For example, the 2-way interactions of *X*_1_*X*_2_ and *X*_3_*X*_4_ on the 4-way combination of *X*_1_*X*_2_*X*_3_*X*_4_, as denoted by [N_4_]_*X*_1_*X*_2_⊥*X*_3_*X*_4__ and calculated as *w*_*X*_1_*X*_2__*w*_*X*_3_*X*_4__ – *w*_*X*_1__*w*_*X*_2__*w*_*X*_3__*w*_*X*_4__, must be accounted for when considering all lower-order effects.

Following the same logic as in the calculation of pairwise interactions, the lower order contribution of drugs, i.e. [N_*N*_]_*D*_1_⊥*D*_2_⊥…⊥*D_n_*_, is given by the expected *N*-way effect when only interactions within the non-single drug subsets (i.e., *D_i_* containing more than one drug) are present. Thus, lower-order effects from the mixture of lower-order combinations and single drugs complementing the total combination will be calculated similarly by replacing the *N*-way drug response, *W*_*X*_1_*X*_2_*X_N_*_, with the product of drug combination responses that are assumed to be only interacting pieces within the *N*-drug combination as

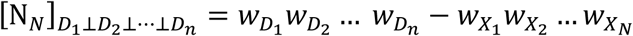

For example the *k*-drug effect of *X*_1_*X*_2_…*X_k_* on the N-way combination for any value of *N*and *k* with *k* < *N* is given by the expected *N*-way effect when only *k* drugs interact, i.e. [N_*N*_]_*X*_1_*X*_2_…*X_k_*⊥*X*_*k*+1_⊥…⊥*X_N_*_ = *W*_*X*_1_*X*_2__…*X_k_ W*_*X*_*k*+1__…*W_X_N__*–*W*_*X*_1__*W*_*X*_2__…*W_X_N__* this is equivalent to the net interaction arising from the *k*-drug combination multiplied by the single drug fitnesses of the remaining *N* – *k* drugs: *W*_*X*_*k*+1__…*W_X_N__*[N_*k*_]_*X*_1_*X*_2_…*X_k_*_.

Subtracting all possible lower-order interaction contributions (via factorizations of *D*_1_ ⊥*D*_2_⊥…⊥ *D_n_*) from the net *N*-way interaction defines newly emergent interactions among all *N* drugs. Accordingly, we now define E_4_ and E_5_ purely in terms of fitness measurements based on the conceptual framework just described and using the above formulas and notation for the lower-order component effects. Guaranteeing that over-counting is eliminated when distinguishing every possible lower-order effect from the net interaction (see S1 Text), the emergent 4-way interaction is given by

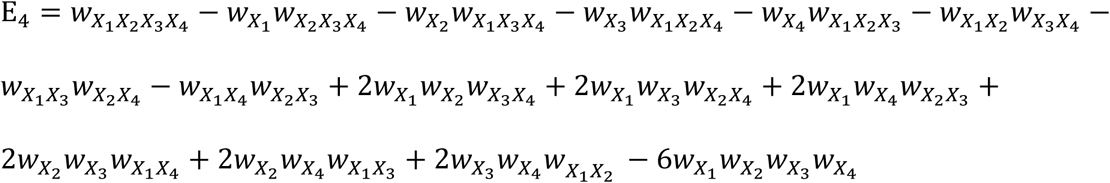

The emergent 5-way measure in terms of relative fitnesses is given as

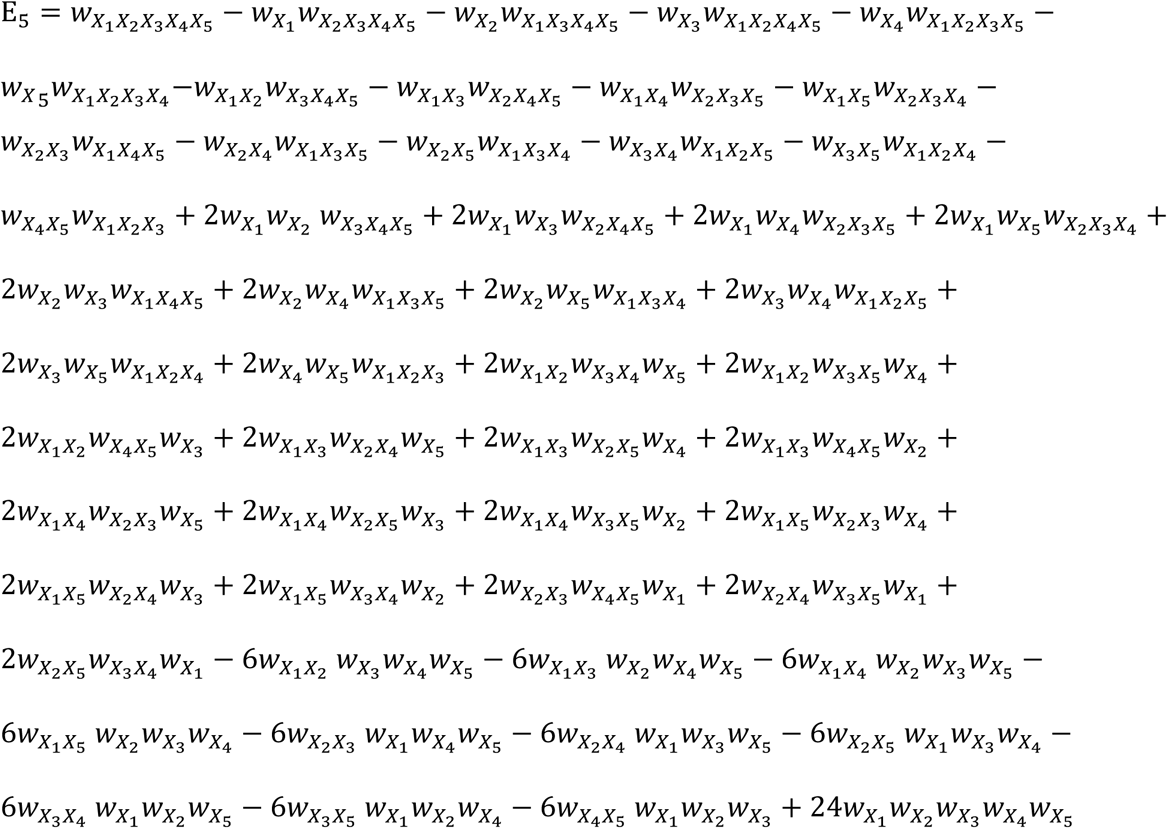

Notably, analyzing deviations from appropriate baselines of these interaction measures allows us to assign information and meaning to the magnitude of the interaction and thus form a correspondence with the type and strength of interaction. This is accomplished via rescaling methods we developed for these interaction measures as defined in^28,29,30^(see S2 Text) and that we use to analyze the experimental data of drug combinations.

#### Details of data analysis and cut-off values for the categorization of interactions

Median growth measurements for each experiment across replicates were used to determine net and emergent interaction types according to the rescaled interaction metric definitions. Due to the full factorial design of our experiments, bacterial growth in the presence of each *k*-drug combination is measured within each drug combination experiment that contains that specific *k*-way drug combination (the pairwise combination *X*_1_*X*_2_ is repeated within the experiments of *X*_1_*X*_2_*X*_3_, *X*_1_*X*_2_*X*_4_, and so on). For such cases (i.e., for 2-, 3-, and 4-drug combinations) the median interaction metric calculation across these experiments was used to determine the interaction type of each drug combination. Given the interaction metric calculation, we categorize synergy, additivity, and antagonism regions according to cut-off values established by previous work ^24, 28, 39^. The interaction among drugs is identified as synergistic when the interaction metric is less than −0.5, antagonistic if it is larger than 0.5, and additive if it ranges between −0.5 and 0.5. Note that the total range of the rescaled metric is from −1 to 1 when the combination of drugs reduces the growth relative to at least one of the lower-order combination growth rates (i.e., fitness), hence leading to rare instances of values above 1.

Finally, we excluded several specific cases in our analysis. First, for 2-drug combinations *X*_4_*X*_2_, the effect of the second drug on the bacterial growth is indistinguishable when both the maximum of single and pairwise growth measurements (i.e., max(*w*_*X*_l__, *w*_*X*_2__) and *w*_*X*_2__) are greater than 90% growth ^40^. Next, using the same reasoning, in the extreme case that *k* drugs by themselves kill off the bacteria populations (lethality: measurements below 4.7% as determined by Tekin et al. ^24^) and the addition of another drug into the environment also leads to lethality, then it is meaningless to look for emergent *k* + 1-drug interactions. We identify these cases as inconclusive and excluded them in our data analysis of the frequency of interaction types.

## Acknowledgements

This work was supported by a James F. McDonnell Complex Systems Scholar Award, an NSF DBI Career award 1254159, an UCLA Faculty Career Development Award, a Hellman Foundation Award, and an NIH/National Center for Advancing Translational Science (NCATS) UCLA CTSI Grant Number UL1TR001881.

## Author contributions

E.T., V.M.S and P.J.Y. conceived of the project and designed the study. C.W. and T.M.K. conducted the experiments, E.T., V.M.S and M.C.L. devised new theoretical analysis tools. R.D. contributed experimental reagents and materials. All authors wrote and revised the manuscript.

